# Hoxa1 and TALE proteins display cross-regulatory interactions and form a combinatorial binding code on Hoxa1 targets

**DOI:** 10.1101/092296

**Authors:** Bony De Kumar, Hugo J. Parker, Ariel Paulson, Mark E. Parrish, Irina Pushel, Brian D. Slaughter, Jay R. Unruh, Julia Zeitlinger, Robb Krumlauf

## Abstract

*Hoxa1* has diverse functional roles in differentiation and development. We have identified and characterized properties of regions bound by Hoxa1 on a genome-wide basis in differentiating mouse ES cells. Hoxa1 bound regions are enriched for clusters of consensus binding motifs for Hox, Pbx and Meis and many display co-occupancy of Pbx and Meis. Pbx and Meis are members of the TALE family and genome-wide analysis of multiple TALE members (Pbx, Meis, TGIF, Prep1 and Prep2) show that nearly all Hoxa1 targets display occupancy of one or more TALE members. The combinatorial binding patterns of TALE proteins defines distinct classes of Hoxa1 targets and indicates a role as cofactors in modulating the specificity of Hox proteins. We also discovered extensive auto- and cross-regulatory interactions among the Hoxa1 and TALE genes. This study provides new insight into a regulatory network involving combinatorial interactions between Hoxa1 and TALE proteins.

## Introduction

Among the clustered *Hox* genes, *Hoxa1* displays the earliest expression during mouse embryogenesis (Hunt et al. 1991; Murphy and Hill 1991) and is one of the most rapidly induced genes during directed differentiation of murine ES cells (Lin et al. 2011; De Kumar et al. 2015). *Hoxa1* has important functional roles in neural crest specification, hindbrain patterning and in heart and ear development (Lufkin et al. 1991; Gavalas et al. 1998; Studer et al. 1998; Gavalas et al. 2001; Makki and Capecchi 2010; Makki and Capecchi 2011; Makki and Capecchi 2012). In clinical studies *Hoxa1* is implicated in etiology and prognosis of various cancers through altered rates of cell proliferation and metastasis (Bitu et al. 2012; Zha et al. 2012; Wardwell-Ozgo et al. 2014; Taminiau et al. 2016). While upstream regulators that establish expression of *Hoxa1* have been characterized (Alexander et al. 2009; Parker et al. 2016), very little is known about its genome-wide binding properties and the downstream target genes through which it exerts it functional roles.

Regulatory analyses of vertebrate *Hox* genes have revealed that they are subject to extensive auto- and cross-regulatory interactions (Tumpel et al. 2009; Parker et al. 2016). The characterization of Hox-response elements associated with these regulatory processes in vertebrate and invertebrate species have uncovered important roles for Pbx/Exd and Meis/Hth as Hox cofactors (Chan et al. 1994; Pöpperl et al. 1995; Rieckhof et al. 1997; Ryoo et al. 1999; Ferretti et al. 2000; Merabet et al. 2007). Pbx and Meis are members of the TALE (<u>T</u>hree <u>a</u>mino-acid <u>l</u>oop extension) family of homeodomain proteins, which consists of six different classes: Irx, Mkx, Meis, Pbx, Prep and TGIF (Burglin 1997). Beyond Pbx and Meis, very little is known about whether there are similar roles for other TALE proteins, as cofactors contributing to Hox binding properties. Pbx and Meis also have more general roles as cofactors for a variety of homeodomain and non-homeodomain classes of transcription factors, (e.g. Pdx1, En, Pax, ZFPIP (Zn finger), PTF1 (bHLH) and T3 receptor-α nuclear hormone receptor). Genetic and regulatory studies have shown that through their roles as general cofactors Pbx and Meis have diverse functional inputs in regulating cell and developmental processes (Moens and Selleri 2006; Laurent et al. 2008; Schulte and Frank 2014).

Studies have characterized physical interactions between Pbx and Hox proteins and shown that these interactions raise their binding specificity and enhance *in vivo* site selection (reviewed in Mann and Chan 1996; Merabet and Mann 2016). This is achieved in part through formation of a Pbx-Hox heterodimer that binds on an overlapping bipartite Hox-Pbx site (Jabet et al. 1999; Piper et al. 1999). Pbx also forms a heterodimer with Meis or Prep using a different domain and this binds to a distinct Pbx-Meis/Prep site (Abu-Shaar et al. 1999; Berthelsen et al. 1999; Penkov et al. 2013). The Hox-Pbx and Pbx-Meis/Prep sites are often in close proximity which facilitates the formation of a ternary complex that helps in fine tuning binding specificity and potentiating regulatory activity (Berthelsen et al. 1998; Ryoo et al. 1999; Ferretti et al. 2000; Ferretti et al. 2005). Meis/Hth proteins have also been shown to be important for controlling the nuclear localization of Pbx/Exd and stability of Pbx mediated complexes (Rieckhof et al. 1997; Abu-Shaar et al. 1999; Berthelsen et al. 1999; Waskiewicz et al. 2001). Pbx/Exd-Hox interactions are involved in repression as well as activation of target genes (Rauskolb and Wieschaus 1994). Hence, TALE proteins as Hox cofactors, may dictate transcriptional regulatory outputs in a context-dependent manner (Galant et al. 2002; Merabet et al. 2003). These studies illustrate the diverse and important roles of cofactors in modulating the binding specificity, selectivity and functional outcomes of Hox proteins.

While there have been extensive studies investigating the conserved roles of Pbx/Exd and Meis/Hth as cofactors for Hox proteins on selected vertebrate and invertebrate loci, there is relatively little information on the contribution of other TALE proteins to Hox specificity. Similarly, very few genome-wide studies have been done in mammalian systems to investigate binding of Hox proteins and their relationships to TALE family members in a tissue-specific or developmental context (Jung et al. 2010; Donaldson et al. 2012; Huang et al. 2012; Sorge et al. 2012). Therefore, the genomewide contribution of TALE proteins to Hox binding specificity is not well understood.

In the current study, we used programmed differentiation of mouse ES cells in combination with genomic approaches to characterize the genome-wide binding properties of Hoxa1 and a set of TALE (Pbx, Meis, Prep1, Prep2, TGIF) proteins. Nearly all Hoxa1-bound regions display occupancy of one or more TALE proteins and their combinatorial binding pattern defines distinct classes of Hoxa1 targets. This shows the TALE family represents a wide repertoire of Hox cofactors. By analyzing Hoxa1 and TALE bound regions and using transgenic reporter assays, we discovered extensive auto- and cross-regulatory interactions among the *Hoxa1* and *TALE* genes. This suggests a *cis*-regulatory mechanism involving feedback loops, by which the expression of Hoxa1 and specific TALE cofactors are co-regulated over time. In summary, this work has generated important new insight into a regulatory network involving combinatorial interactions between Hoxa1 and TALE proteins that defines the binding specificity of Hoxa1 on its target genes.

## Results

### Characterization of ES cell line carrying an inducible epitope-tagged variant of Hoxa1

In this study, we sought to characterize the genome-wide occupancy of Hoxa1 and to understand the nature of its downstream targets. We used the programmed differentiation of ES cells into neuroectoderm as a model system, due to the transient expression of *Hoxa1* in development and limiting amounts of appropriate embryonic neural tissue. A variety of studies have shown that human and mouse ES cells can be differentiated into neuronal fates using retinoic acid (RA) and Hox genes are sequentially activated in a manner that mimics their induction during embryonic development (De Kumar et al., 2015; Mazzoni et al., 2013; Papalopulu et al., 1991; Sheikh et al., 2014; Simeone et al., 1990; Simeone et al., 1991). Previously we have extensively characterized RA-induced differentiation of KH2 ES cells and provided insight into the mechanistic basis for rapid and early induction of *Hoxa1* in ES cells and mouse embryos through paused polymerase and control of transcriptional elongation (Lin et al. 2011; Gaertner et al. 2012; De Kumar et al. 2015). In the absence of a validated Hoxa1 antibody for genomic studies, we generated a KH2 ES cell line (Beard et al. 2006) with a locus specific insertion (Col 1A) carrying a doxycycline (Dox) inducible version of Hoxa1 tagged with 3X-Flag and Myc epitopes at the C-terminus (Fig. 1A). We optimized the conditions to ensure that the expression levels of the epitope-tagged version of *Hoxa1* in response to RA were comparable to the endogenous gene. By quantifying levels using three independent methods: singlemolecule RNA fluorescent *in situ* hybridization (FISH), and qPCR and RNA-seq we found that the epitope tagged version was expressed in a ratio of 1:1.5 compared to endogenous Hoxa1(Supplemental Fig. S1B-D). Furthermore, we analyzed the time course of expression of the tagged Hoxa1 protein by western hybridization and found that it is detected as early as 6 hrs following induction and levels increase up to 48 hrs (Supplemental Fig. S1A).

**Figure 1.**
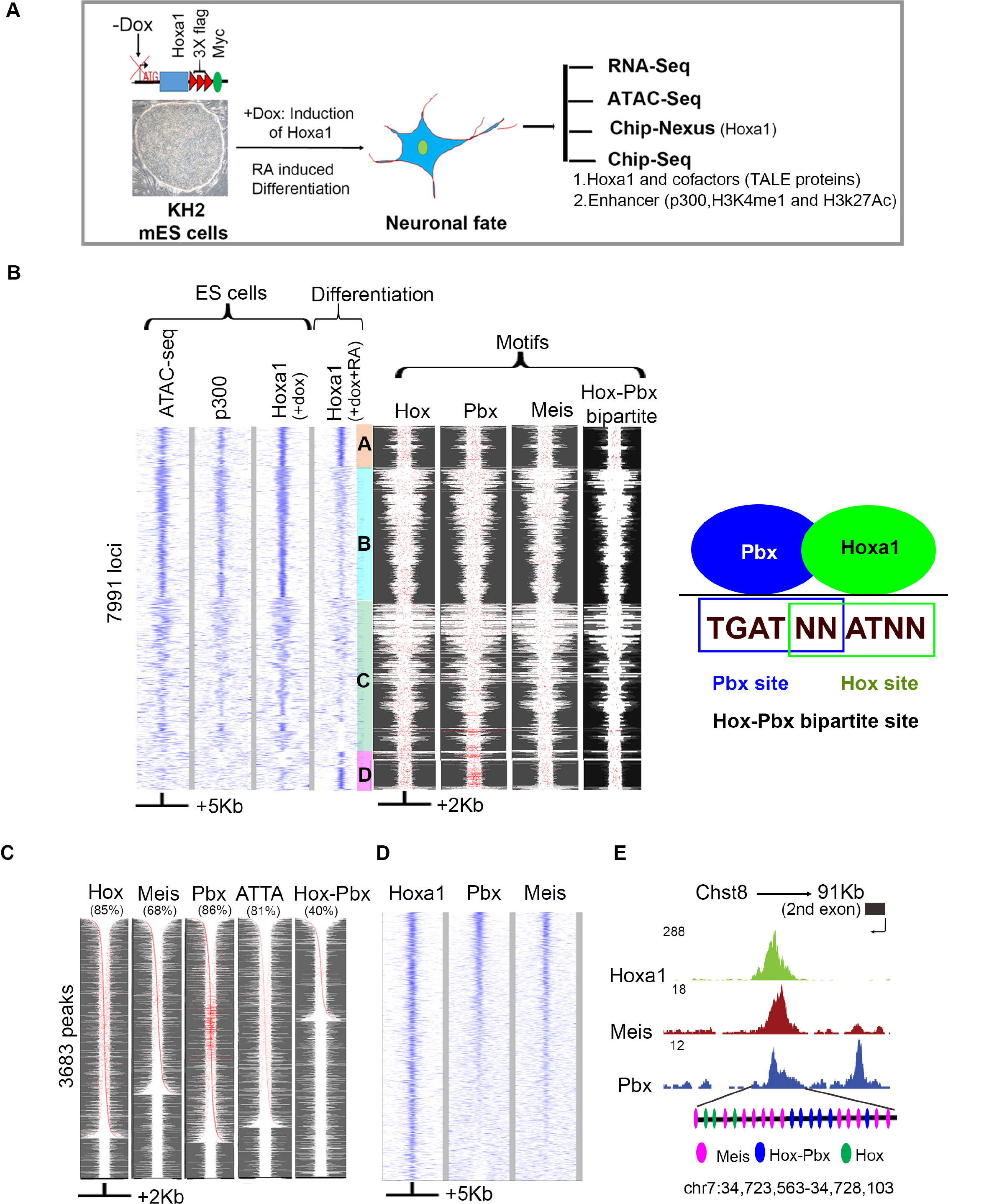
Genome-wide identification and analysis of Hoxa1 bound regions reveals overlaps with Pbx and Meis binding. (**A**) Outline of experimental design for differentiation and analysis. Addition of Dox (+Dox) induces epitope tagged *Hoxa1* and addition of retinoic acid (+RA) induces programmed differentiation of ES cells into neural fates. (**B**) Heatmaps of genome-wide occupancy of Hoxa1 in ES cells (+dox) and differentiated cells (+dox +RA) defined by ChIP-seq. The occupancy of p300 and chromatin state (ATAC-seq) in ES cells is also shown. Comparative analyses between ES and differentiated cells reveals four distinct groups (A-D) of bound regions. The four columns on the right, show the spatial distribution of consensus motifs for Hox, Pbx, Meis and Hox-Pbx bipartite binding sites in the corresponding genomic regions. All variants of the Hox, Pbx and Meis motifs in the TransFac database along with a consensus motif for Hox-Pbx bipartite sites (TGATNNATNN) were searched for using FIMO. Best matches (p<=1e-4) are shown in red. At the far right is a diagram illustrating the relationship of Pbx and Hox binding sites in Hox-Pbx bipartite motifs. Groups A and D, representing Hoxa1 bound regions in differentiated cells showed increased presence of Hox-Pbx bipartite sites and tandem clusters of Pbx motifs. **(C**) Hoxa1 bound regions in differentiated cells are enriched for a variety of consensus Hox, Pbx and Meis motifs. For each TransFac motif, FIMO matches (p<=1e-4) are shown in red. The respective motif and *%* of peaks with the motif are indicated at the top of each column. Rows are in different orders for each column, and are sorted by the distance of the central-most motif to the peak center. Many Hoxa1 bound regions in a given row show multiple red dots indicative of multiple binding motifs in a peak. (**D**) Heatmap shows extensive co-occupancy of Hox, Pbx and Meis on Hoxa1 targets in differentiated cells. The plots use IP coverage of z-score matrix for Pbx and Meis binding at Hoxa1 peaks. Rows are sorted by the intensity of the Pbx signal within the central 1kb region. In **B-D,** the size of regions flanking the center of the peak of Hoxa1 binding are shown at the bottom. (**E**) UCSC genome browser shots showing Pbx, Meis, and Hoxa1 in exon 2 of *Chst8*. Clustering of multiple Hox, Hox-Pbx and Meis binding sites are also shown at the bottom for the Hoxa1 bound region. Genomic coordinates (mm10) are indicated at the bottom

### Clustered binding sites for Hox, Pbx and Meis

Using the Dox-inducible epitope-tagged Hoxa1 ES cell line, we then performed ChlP-seq experiments. The induction of Hoxa1 by Dox in the absence of RA treatment, identified 8836 genomic regions near 6029 genes that were bound by Hoxa1 (Fig.1B, Supplemental Table S1). These peaks are found in regions of open chromatin, as assayed using ATAC-seq (Assay for Transposase-Accessible Chromatin with high throughput sequencing) and show occupancy of co-activators (p300) (Fig. 1B). In contrast, there is a marked change in the binding profile upon RA-induced differentiation. A subset of the Hoxa1 bound regions in ES cells retain occupancy upon differentiation (group A), while the majority of the other regions showed a reduction or loss of binding (groups B and C) (Fig. 1B), leaving a total of 3682 Hoxa1-bound regions near 2595 genes in differentiated cells (Supplemental Table S2). However, there is a new group of binding sites in differentiated cells (group D), which were not found in previously open chromatin and did not display binding of p300 in ES cells (Fig. 1B). Hoxa1 may function as a pioneer factor on these new target sites.

We compared the properties of these bound regions from undifferentiated and differentiated ES cells and found that the peaks in ES cells show a relatively low fold enrichment over input, clustered around 2-fold. While in differentiated cells, the major fraction of Hoxa1 bound peaks show a higher degree of enrichment (3-6 fold) (Supplemental Fig. S1A). By ranking peaks from both samples based on fold enrichment, nearly all top ranking peaks correspond to bound regions in differentiated cells (Supplemental Fig. S1B). This suggests that the major proportion of Hoxa1 bound regions in ES cells, have a lower occupancy compared to sites bound after differentiation. This difference does not correlate with the number of peaks per gene as these were similar in both samples. We next analyzed motif content and searched for differences in enriched consensus binding sites using HOMER. This analysis revealed that both Hox and Hox-Pbx bipartite sites are significantly enriched in regions occupied by Hoxa1 in differentiated cells (Supplemental Fig. S1C and Supplemental Table S3,4). This suggests that the increase in occupancy in differentiated cells may be mediated in part by the association of Hoxa1 with Pbx and Meis on Hox-Pbx bipartite sites.

Beyond Hox-Pbx bipartite sites, Pbx, Meis and Pbx/Meis heterodimers also bind a variety of unique and overlapping consensus motifs (Penkov et al. 2013). Hence, we extended our analysis to search for the presence and spatial distribution of these motifs in Hoxa1 bound regions in ES cells versus differentiated cells using FIMO and motif definitions in the TransFac database (Fig. 1B, C). The subset of Hoxa1 bound regions that retained occupancy upon differentiation (group A) and the set of new binding sites that appeared in differentiated cells (group D), both showed increased presence of Hox-Pbx bipartite sites and tandem clusters of Pbx motifs (Fig. 1B). This supports the hypothesis that the differential binding properties of Hoxa1 in these two cell states is related to differences in the *cis*-regulatory motif composition.

Focusing on the peaks bound in differentiated cells, they contain known consensus motifs for Hox (85%), Pbx (86%), Meis (68%) and Hox-Pbx bipartite sites (40%) which are often present in multiple copies per peak (Fig. 1C). This raises the question of whether Pbx and Meis are physically occupied on these genomic loci along with Hoxa1. To investigate occupancy of Pbx and Meis, we performed ChIP-Seq experiments using α-Pbx1/2/3 and α-Meis1/2 antibodies. We identified 5761 Pbx and 1410 Meis binding regions (1% IDR) enriched for consensus motifs that correlate with those previously identified in 11.5 day mouse embryos (Penkov et al. 2013) (Supplemental Table S5,6). The larger number of Pbx bound regions compared to Meis is consistent with its more general role as a cofactor for a variety of TFs (Laurent et al. 2008; Schulte and Frank 2014). In comparing these genome-wide profiles with Hoxa1 bound regions, a large number also display co-occupancy of Pbx and Meis (Fig. 1D). This is illustrated by Hoxa1 binding at the *Chst8* locus, where multiple Meis, Hox and Hox-Pbx bipartite sites are present and there is evidence for physical occupancy of these transcription factors (Fig. 1E).

To examine Hoxa1 binding at high-resolution we used a ChIP-nexus approach (He et al. 2015). This is based on the use of lambda exonuclease during ChIP to determine the exact location where Hoxa1 is cross-linked. The ChIP-nexus data revealed that single peaks identified by ChIP-seq often represent clusters of multiple binding events. HOMER analysis of the ChIP-nexus binding profiles showed that Hoxa1 was not only bound to Hox and Hox-Pbx bipartite motifs but also to Pbx and Meis consensus motifs (Supplemental Table S7). This implies that Hoxa1 may bind to DNA directly or indirectly through association with Pbx and Meis as cofactors. Together, this data indicates that on a genome-wide basis, Hoxa1 binding is frequently associated with Pbx and Meis proteins and that a wide variety of motif combinations underlies the binding of Hoxa1.

### Distinct combinations of TALE proteins on Hoxa1 bound regions

Pbx and Meis are not the only members of the TALE family of homeodomain proteins, which also includes TGIF, Prep1, Prep2, Irx and Msx (Burglin 1997; Mukherjee and Burglin 2007; Longobardi et al. 2014). Since Hoxa1-bound regions are frequently co-occupied by Pbx and Meis, we investigated whether other TALE proteins may also be associated with Hoxa1 bound regions. In ES cells Pbx1, TGIF and Prep2 are expressed at low levels, but upon RA treatment these and other TALE cofactors are induced and become available to partner with Hoxa1 (Supplemental Fig. S3). We therefore performed additional ChIP-seq experiments with α-TGIF, α-Prep 1 and α-Prep2 antibodies to compare the genome-wide binding profiles of the TALE family in both ES cells and differentiated cells.

Our analysis revealed that nearly all Hoxa1 bound regions display occupancy of one or more TALE proteins, yet each TALE protein has a unique binding pattern (Fig. 2). Eight classes were identified through k-means clustering (Cluster 1-8), based on their distinct relative levels of TALE proteins and their spatial relationship to the center of Hoxa1 binding (Fig. 2A, B). Analysis of the *Fgfr1* and *Zfp703* loci provide specific examples of distinct classes of binding along with their epigenetic states and enhancer properties (Fig. 2C, D).

**Figure 2.**
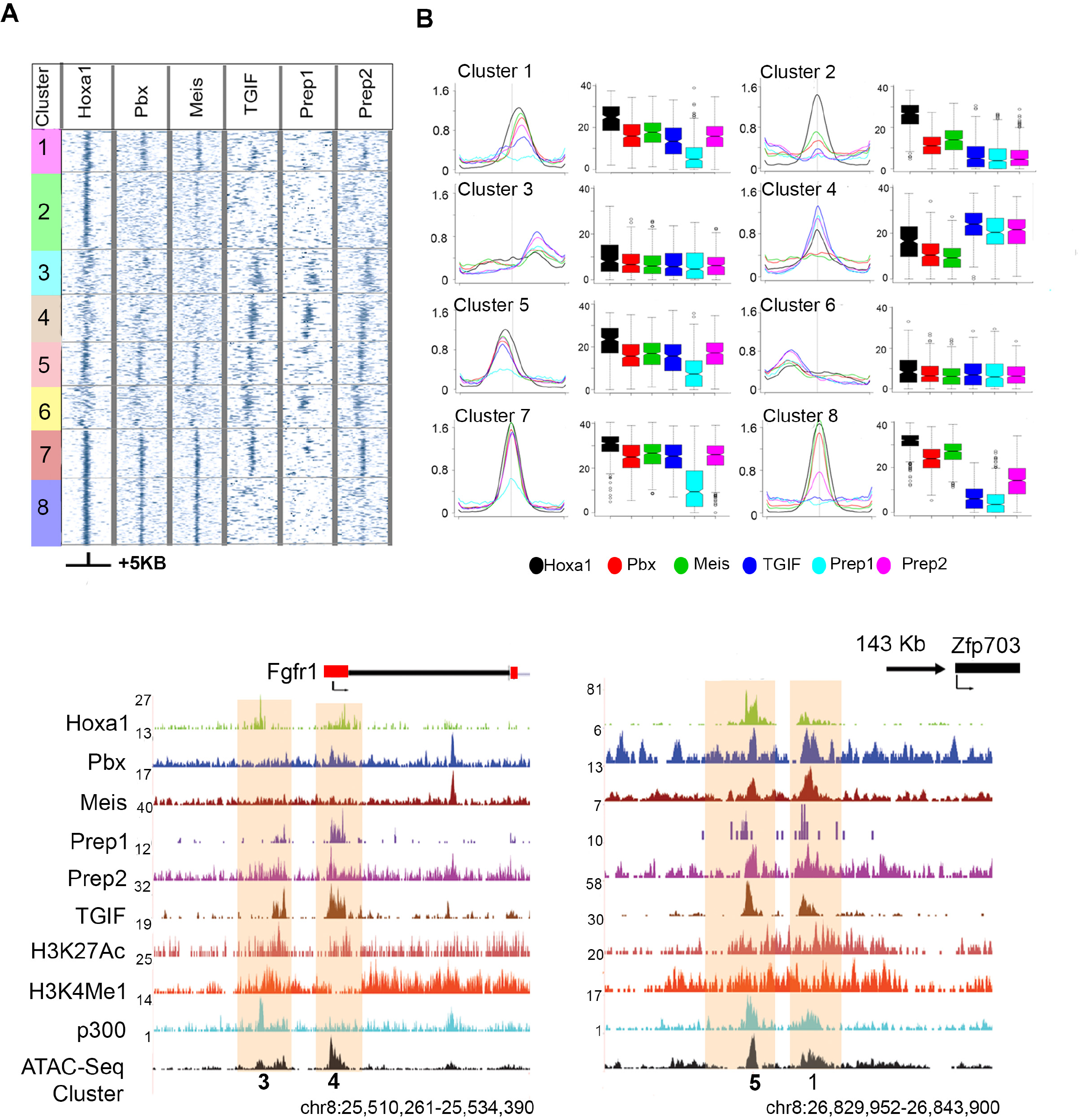
Patterns of co-occupancy of TALE proteins on Hoxa1 targets define 8 distinct clusters. (**A**) Heatmaps for occupancy of individual TALE factors at Hoxa1 peaks using IP coverage of z-score matrix after kmeans-clustering with k=8. Only positions of positive z-score are shown. A +/- 5kb region flanking the center of the peaks of Hoxa1 binding are shown. (**B**) Average binding levels, shown as line and Box plots, illustrate the distinct properties with respect to relative binding of each protein and its spatial distribution of each cluster in **A**. Boxplots are row sums from the central 1 kb of each column. (C) UCSC browser shots (mm10) of *Fgfr1* and *Zfp703* loci, showing Hoxa1 bound regions along with occupancy of TALE cofactors, activator protein p300, enhancer marks and chromatin state. The clusters to which these bound regions map are noted at the bottom.

The characteristics of these classes uncover distinct and novel interactions of TALE family members on Hoxa1 bound regions. Cluster 8 represents a classical group in which Pbx and Meis are the primary co-factors for Hoxa1, with little or no binding of other TALE members. In contrast, Cluster 4 illustrates a group of targets with little or no occupancy of Pbx and Meis, but high levels of binding by the other TALE proteins. Cluster 7 illustrates a class where nearly all TALE proteins co-occur (Fig. 2).

To explore whether the different clusters of target genes are associated with different functions, we performed a KEGG pathway analysis on the genes associated with each cluster. The results show an enrichment of distinct and overlapping pathways for each cluster (Supplemental Fig. S4). For example, Clusters 3, 4, 6 and 8 are all enriched for components of the Hippo signaling pathway. Clusters 4 and 6 are also enriched for Hedgehog signaling, while Clusters 6 and 8 are enriched for Wnt signaling. This suggests that the binding patterns observed in different clusters may be functionally relevant for context-dependent regulation of diverse biological processes by Hoxa1. Together, these findings imply that the TALE family provides a wide repertoire of Hox cofactors beyond Pbx and Meis.

### TALE binding preferences are associated with distinct classes of *cis*-motifs

The clusters of differential occupancy of TALE proteins raises the possibility that in each cluster there is a distinct combination of motif signatures at the level of *cis*-elements (combinatorial code) which generates permutations that mediate differential recruitment of TALE proteins to the Hoxa1-bound regions. To explore the basis of differential binding of TALE proteins on Hoxa1 targets, we first analyzed the general motif composition of the genome-wide binding preferences of individual TALE proteins. This generated a series of known and novel enriched consensus motifs for each protein. We then compared the relative enrichment of these motifs in regions bound by other TALE proteins (Supplemental Table S6, 8). In this manner, we obtained nine distinct classes of motif signatures that reflect unique and overlapping combinations of binding preferences (Fig. 3A, Supplemental Table S8). For example, motifs M1-M4 are enriched for all TALE proteins, while M25-M34 are specifically enriched in Prep1 bound regions.

**Figure 3.**
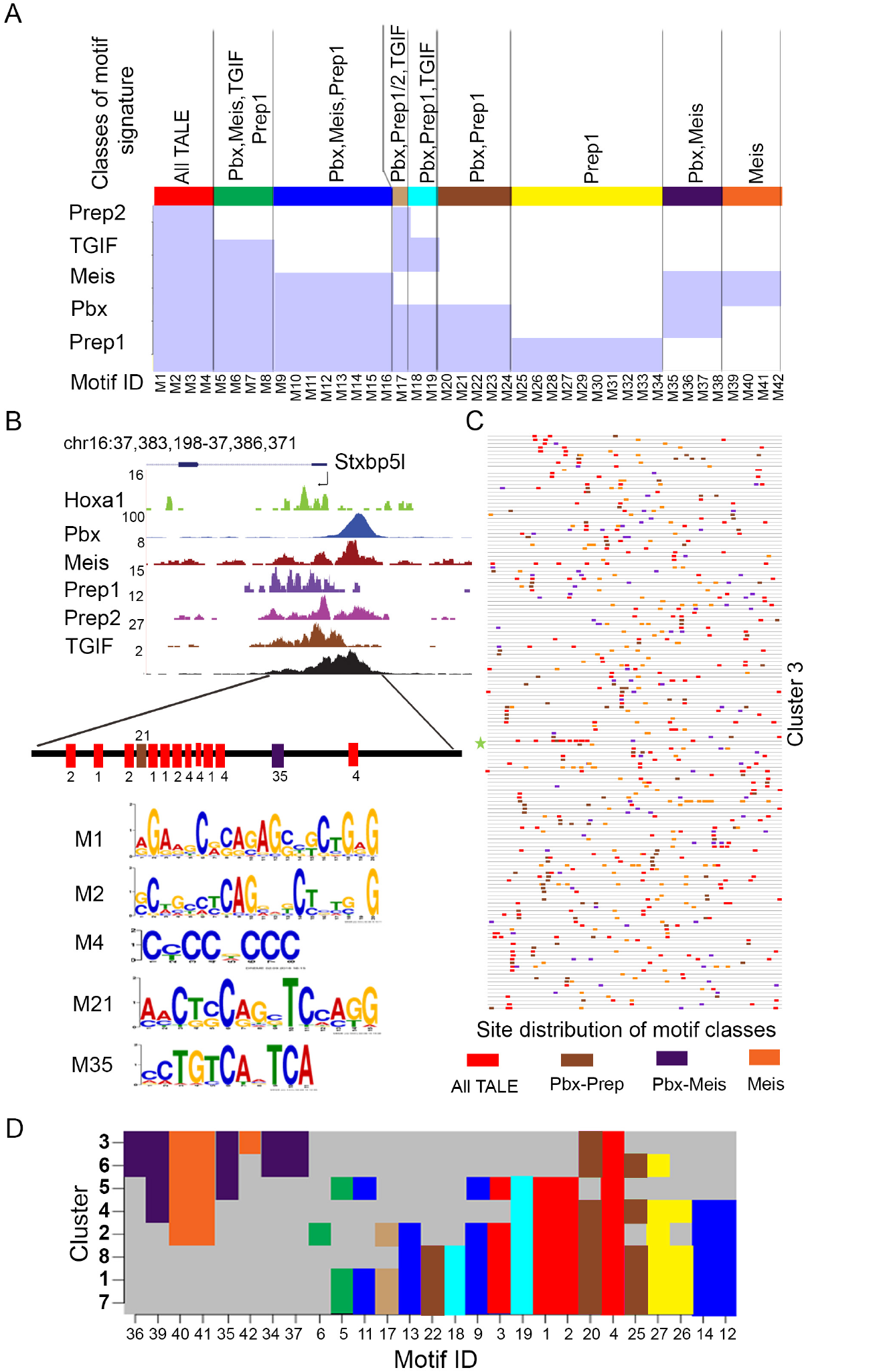
Unique and overlapping genome-wide binding preferences of individual TALE proteins and their distribution in Hoxa1 bound regions. **(A)** Analysis of enriched motifs from genome-wide binding preferences for members of the TALE family (Pbx, Meis, TGIF, Prep1 and Prep2) using Meme and AME. There are 42 enriched motif signatures (M1-M42) comprised of both novel and known consensus TALE binding sites. These 42 *cis*-motifs are divided into nine distinct classes based on their presence or absence within regions bound by one or more of the TALE family members, as indicated by the shaded light blue areas in the matrix. At the top, each class is identified by a separate color along with the respective TALE family members showing occupancy on these motifs. **(B)** UCSC browser shot showing occupancy of Hoxa1 and TALE factors near the *Stxbp5l* locus which is a representative binding region in Cluster 3. The diagram at the bottom indicates that it contains 5 different motifs (M1, M2, M4, M21 and M35) representing 4 classes of motif signatures defined in (**A)** above. There are multiple copies of 3 of the *cis*-motifs (M1, M2 and M4). Genomic coordinates (mm10) are indicated at the top. **(C)** A block diagram shows the occurrence of four distinct motif classes with multiple copies in all the loci present in Cluster 3. Each respective motif class is shown as a distinct color using those indicated in (**A)**. Length of peaks are normalized to a standard length for uniformity of display. (**D**). A matrix indicating the presence or absence of the individual classes of cis-motif signatures in the respective clusters defined by combinatorial TALE binding on Hoxa1 bound regions. Distribution various motif signature classes in each cluster are shown in color to match those in **A**. This plot indicates unique combinations of motif signatures which underlie the differential binding of TALE proteins in each cluster.

We then looked for the presence or absence of these individual classes of *cis*- motif signatures in the respective clusters defined by TALE binding and found that with the exception of Clusters 1 and 7, each cluster had a distinct and unique combination of motif signatures (Fig. 3D). Hence, the binding patterns found in Clusters 1-8 correlate with the underlying specific patterns of motif enrichment. For example, in Cluster 5, Prep1 displays a low level of occupancy compared to the other TALE proteins, and there is a complete absence of Prep1 specific motifs (Fig. 2, 3D). Cluster 2-6 are enriched for Meis/Pbx and Meis classes and have a reduced repertoire of other classes of motifs compared to the other clusters. Together, this data suggests a mechanistic basis via a combinatorial *cis*-binding code, for differential recruitment of TALE proteins to the Hoxa1-bound regions.

In general, the clusters of differential TALE occupancy in Hoxa1 bound regions consists of multiple *cis*-motifs from these classes. *Stxbp5l* is an example of a binding profile of a locus from Cluster 3 containing 5 different motifs, representing 4 classes of motif signatures (Fig. 3B). There are multiple copies of 3 of the cis-motifs (M1, M2 and M4). Similar patterns with multiple motifs and classes are observed for all loci in Cluster 3 (Fig. 3C). These findings further reinforce the observation that in Hoxa1 bound regions there are distinct combinations of multiple binding motifs for both Hoxa1 and TALE proteins. Analysis using HOMER, identified enriched motifs for other factors in each cluster that may also contribute to differential patterns of binding (Supplemental Table S9).

### Auto- and cross-regulatory interactions between Hoxa1 and TALE genes

In examining the putative target genes of Hoxa1 and TALE proteins, we found evidence for extensive occupancy of these proteins on the genes that encode them, suggesting cross-regulatory interactions between the regulators themselves. We therefore analyzed how such regulatory network might function during ES cell differentiation. Since Pbx, TGIF and Prep2 are expressed in ES cells at low levels, we performed ChIP-seq experiments for these factors in ES cells. We found that the *Hoxa1* gene itself is a downstream target, as a region upstream of the Hoxa1 promoter is bound by Pbx, TGIF and Prep2 in ES cells. This is illustrated by a browser shot of the *Hoxa1* locus and the combined regulatory inputs are depicted as a Biotapestry model (Longabaugh et al. 2005) (Fig. 4A). The region is accessible but epigenetic enhancer marks are absent. This is consistent with the observation that while inactive in ES cells, paused polymerase is loaded on the *Hoxa1* promoter (Lin et al. 2011; De Kumar et al. 2015).

**Figure 4.**
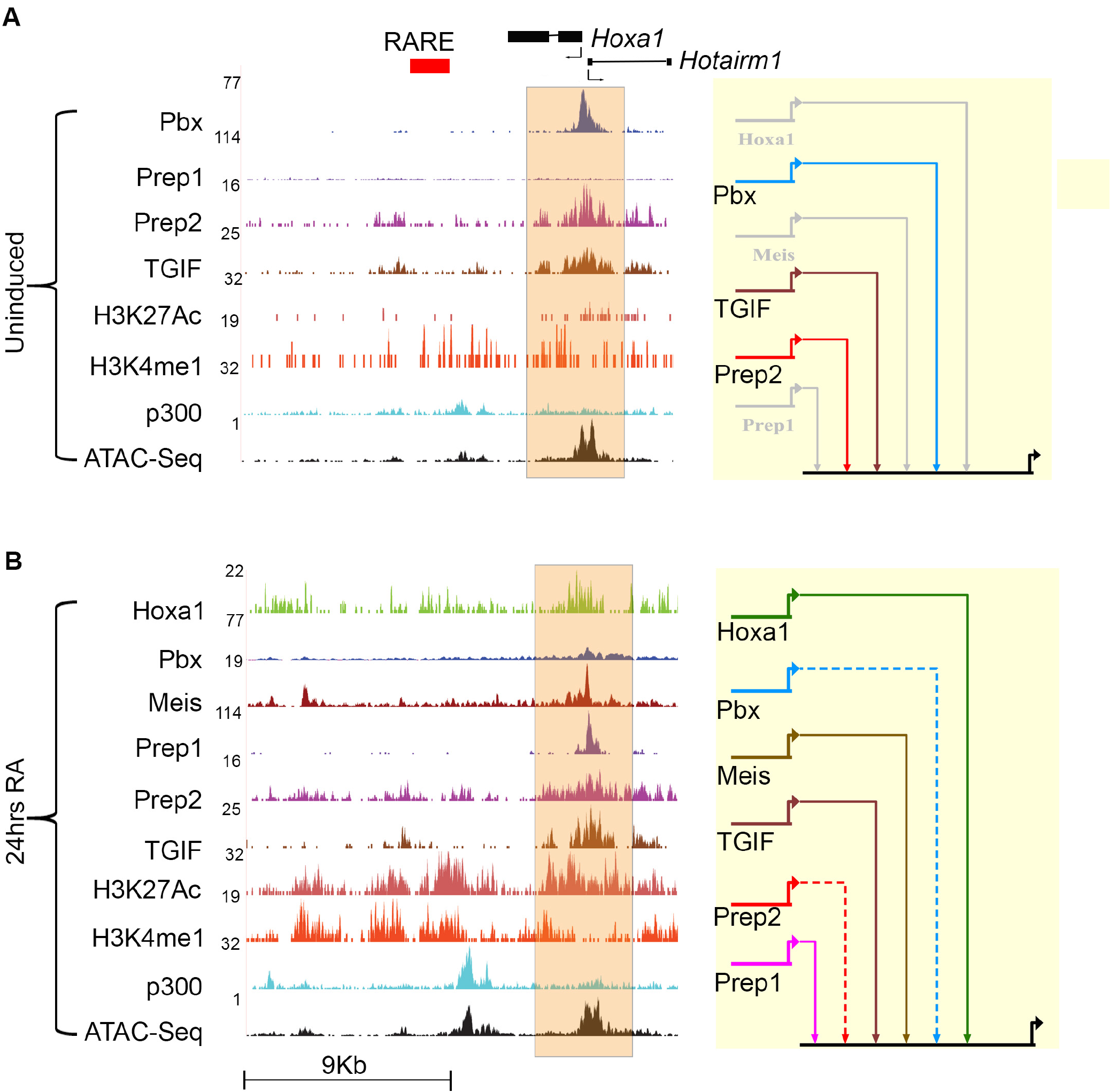
Hoxa1 and TALE proteins bind to the *Hoxa1* locus. (**A, B**) UCSC genome browser shots (mm10) for the *Hoxa1* locus showing binding of Hoxa1 and occupancy of the TALE family members in ES cells (**A**) and differentiated (**B**) cells. The presence of modified histone marks (H3K4me1 and H3K27Ac) characteristic of enhancers, occupancy of the p300 activator protein and accessibility of chromatin (ATAC-Seq) at these genomic loci are also shown. The shaded area indicates *Hoxa1* bound regions near the promoter. In the Biotapestry plots at the right of each panel, the solid colored line indicates binding of each specific TALE factor, while grey lines indicate lack of binding in the given cellular states. Dotted line represent decrease in binding over regions.

During the rapid induction of Hoxa1 expression upon RA induced differentiation the occupancy of TALE proteins and their epigenetic state undergo a dynamic change (Fig. 4B). Pbx and Prep2 binding are reduced, while Meis and Prep1 are recruited to this site along with the appearance of histone modifications associated with active enhancers. In addition, we observe increased occupancy of Hoxa1 over this region, suggesting that it provides an auto-regulatory input into its own expression (Fig. 4B).

In an analogous manner, we find that the genes encoding TALE proteins are themselves downstream targets of Hoxa1 and TALE cofactors (Fig. 5, Supplemental Fig. S5). This data suggests the presence of extensive auto- and cross-regulatory interactions among TALE genes and that Hoxa1 plays a role in regulating the TALE cofactor genes. To test whether these auto- and cross-regulatory elements are functional *in vivo*, we used reporter assays in zebrafish to monitor the activity of 13 of these regions. We found that 11 are able to mediate reporter expression in neural tissues (Fig. 5-7 and Supplemental Fig. S5). For example, TALE and Hoxa1 bound regions flanking *Meis1, Pbx1* and *TGIF* mediate expression of a GFP reporter in the nervous system (Supplemental Fig. S5). This suggests that they may function as enhancers contributing to regulation of TALE genes.

**Figure 5.**
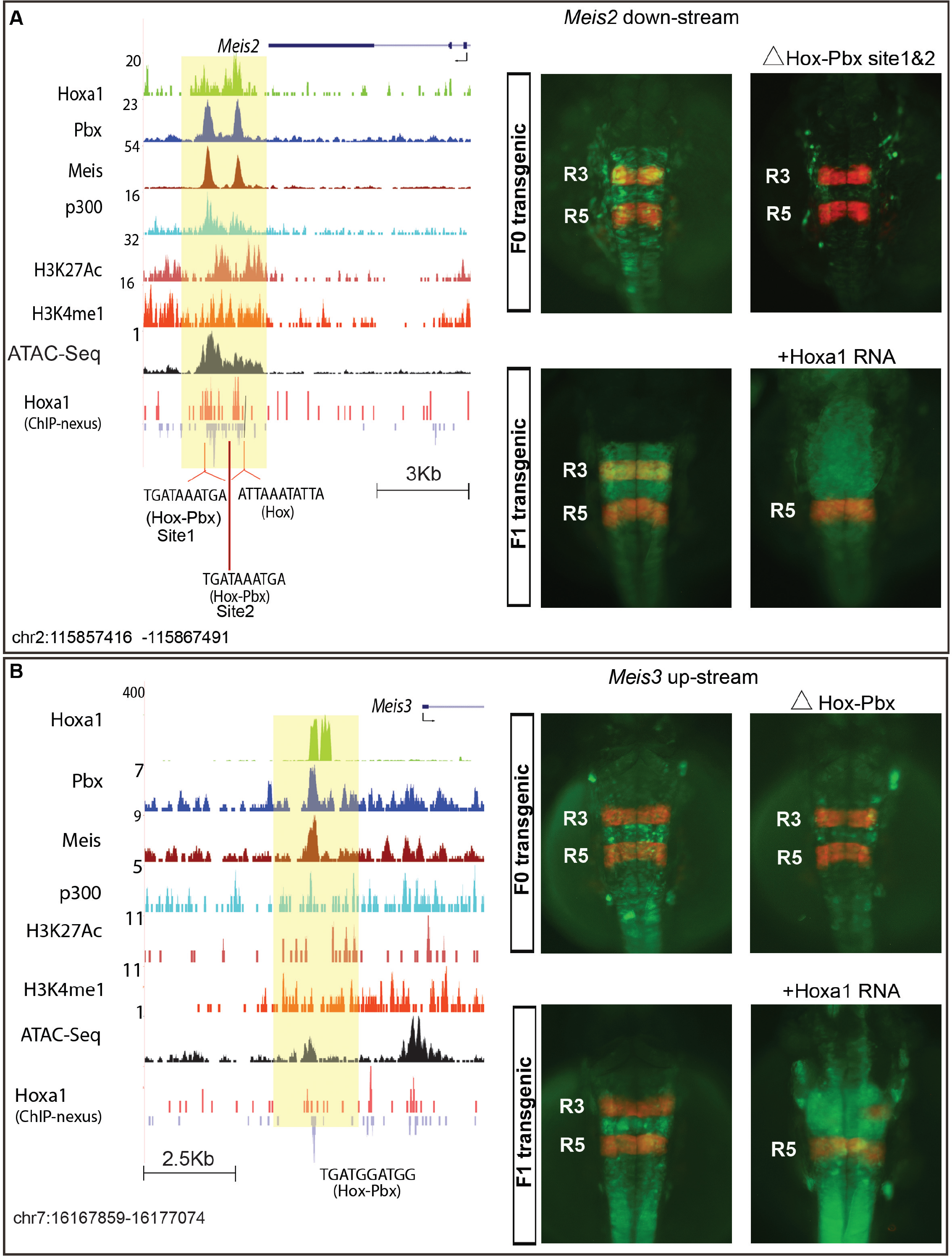
The *Meis2* and *Meis3* genes are down-stream targets of Hoxa1. (**A, B**) UCSC browser shots of *Meis2* (**A**) and *Meis3* (**B**) showing Hoxa1 bound regions along with occupancy of Pbx and Meis cofactors, activator protein p300 and open chromatin states are shown. ChIP-nexus data reveals the presence of multiple binding motifs for Hoxa1 and its cofactors in the protected regions as noted at the bottom. The shaded area indicates bound regions. Genomic coordinates (mm10) are indicated below each locus. Adjacent to each browser shot at the right are transgenic reporter assays testing the regulatory potential of the binding regions +/- 250 bp. For both *Meis2* and *Meis3*, F_0_ and F_1_ transgenic embryos show restricted reporter expression in the hindbrain mediated by the wild type fragments. The wild type enhancers respond to ectopic expression of *Hoxa1* as indicated by the loss of RFP (red) specific expression in R3 and expansion of GFP (green) expression. Deletion of the Hox-Pbx sites in these regions abolishes GFP reporter activity. This demonstrates that the binding regions function as Hox-dependent response elements *in vivo*.

While we identified various motif combinations among Hoxa1-bound regions, we noticed that among the validated enhancers many were consensus Hox-Pbx bipartite sites. These sites showed specific footprints in ChIP-nexus data (Fig. 5 and Supplemental Fig. S5). To functionally validate the role of this class of sites, we assayed cis-regulatory regions from *Meis2* and *Meis3*, which both had enhancer related histone modifications and Hox-Pbx bipartite motifs (Fig. 5A, B). Upstream of *Meis3* there is a single Hox-Pbx site, while in the case of *Meis2*, a 3’ flanking region contains two Hox-Pbx motifs (Site1 & Site2) and a Hox site. However, no binding was detected on Hox-Pbx Site2. Using transgenic reporter assays, we found that both transient transgenic (F_0_) and F_1_ lines of zebrafish carrying both of these regions are able to direct GFP reporter expression in the embryonic hindbrain (Fig. 5A, B). Specific deletion of the Hox-Pbx motifs from these regions leads to the marked reduction of transgene expression in both cases, showing that the motifs are required for proper regulatory activities of the enhancers. In the *Meis2* locus, mutation of Hox-Pbx Site2, which lacked binding of Hoxa1, had no effect on transgene activity (data not shown). To test whether these motifs are responsive to *Hoxa1* dosage, we assayed the reporter expression after injection of *Hoxa1* mRNA. Indeed, the expression domains of the *Meis2* and *Meis3* reporters expanded into more anterior regions, indicating that they are responsive to *Hoxa1 in vivo* (Fig. 5A, B). This shows that the Hox-Pbx bipartite sites, bound by Hoxa1, are indeed functional *in vivo* and implies that these Hox-Pbx bipartite sites play important roles in potentiating *Hoxa1*-mediated regulation of TALE genes. From a gene regulatory network perspective, these interactions imply that *Hoxa1* utilizes a feedforward regulatory circuit to control the expression of TALE cofactors important for its function during patterning and development (Fig. 6B).

**Figure 6.**
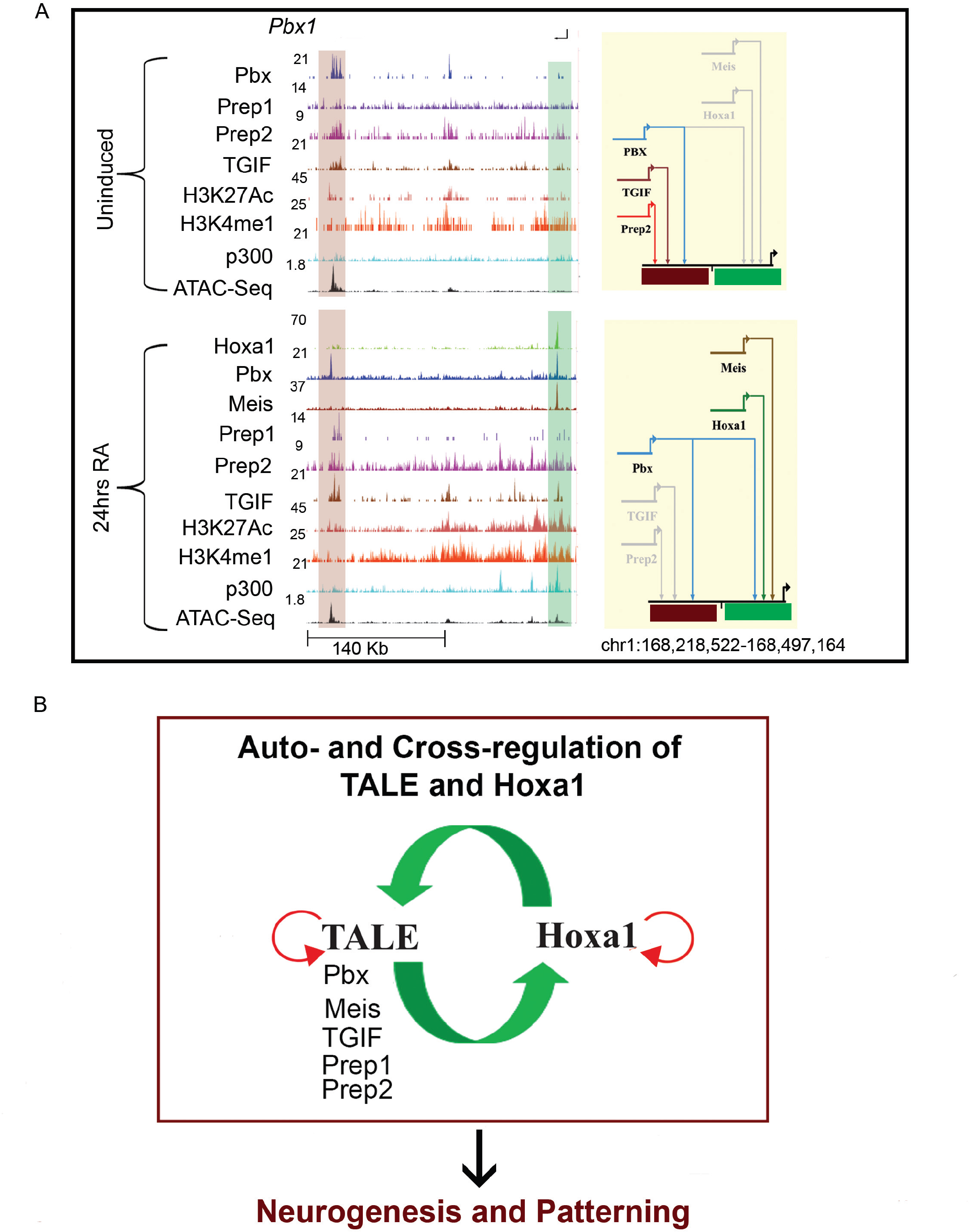
Hoxa1 and TALE proteins bind to the *Pbx1*. (**A,**) UCSC genome browser shots (left) for the *Pbx1* locus showing binding of Hoxa1 and occupancy of the TALE family members in ES cells (top) and differentiated cells (bottom). The presence of modified histone marks (H3K4me1 and H3K27Ac) characteristic of enhancers, occupancy of the p300 activator protein and accessibility of chromatin (ATAC-Seq) at these genomic loci are also shown. The shaded area indicates bound regions. In the Biotapestry plots at the right of each panel, the solid colored line indicates binding of each specific TALE factor, while grey lines indicate lack of binding in the given cellular states. Genomic coordinates (mm10) are indicated below each locus. (**B**) Model summarizing auto- and cross-regulatory interactions among *Hoxa1* and *TALE* genes and their output on downstream targets.

Mapping the diverse inputs of Hoxa1 and TALE proteins into cis-regulatory regions adjacent to TALE genes indicates a complex array of cross-regulatory modules. There are diverse patterns of binding on multiple sites spread in and around these loci and we observed changes in occupancy during differentiation (Fig. 6A and 7). For example, *Pbx2* has a single 5’ cis-regulatory element that changes its occupancy during differentiation. In ES cells, this region is occupied by Prep2 and Pbx, but upon differentiation these factors are replaced by occupancy of Hoxa1, Prep1 and TGIF. Likewise, there are two regions flanking the Pbx1 gene, which are bound by Pbx, TGIF and Prep2 in ES cells but this binding pattern is lost upon RA treatment (Fig. 6A and 7). Instead, Hoxa1, Meis and Pbx are recruited to a second distinct downstream region upon RA treatment.

These extensive cross-regulatory inputs amongst TALE genes and Hoxa1 can be visualized using Biotapestry based topological models (Fig. 7). These models indicate that in most cases, an array of cis-elements integrates dynamic inputs by Hoxa1 and TALE proteins to modulate their expression during differentiation. These interactions contribute to the availability and levels of TALE proteins which impact Hoxa1 binding on its downstream targets.

**Figure 7.**
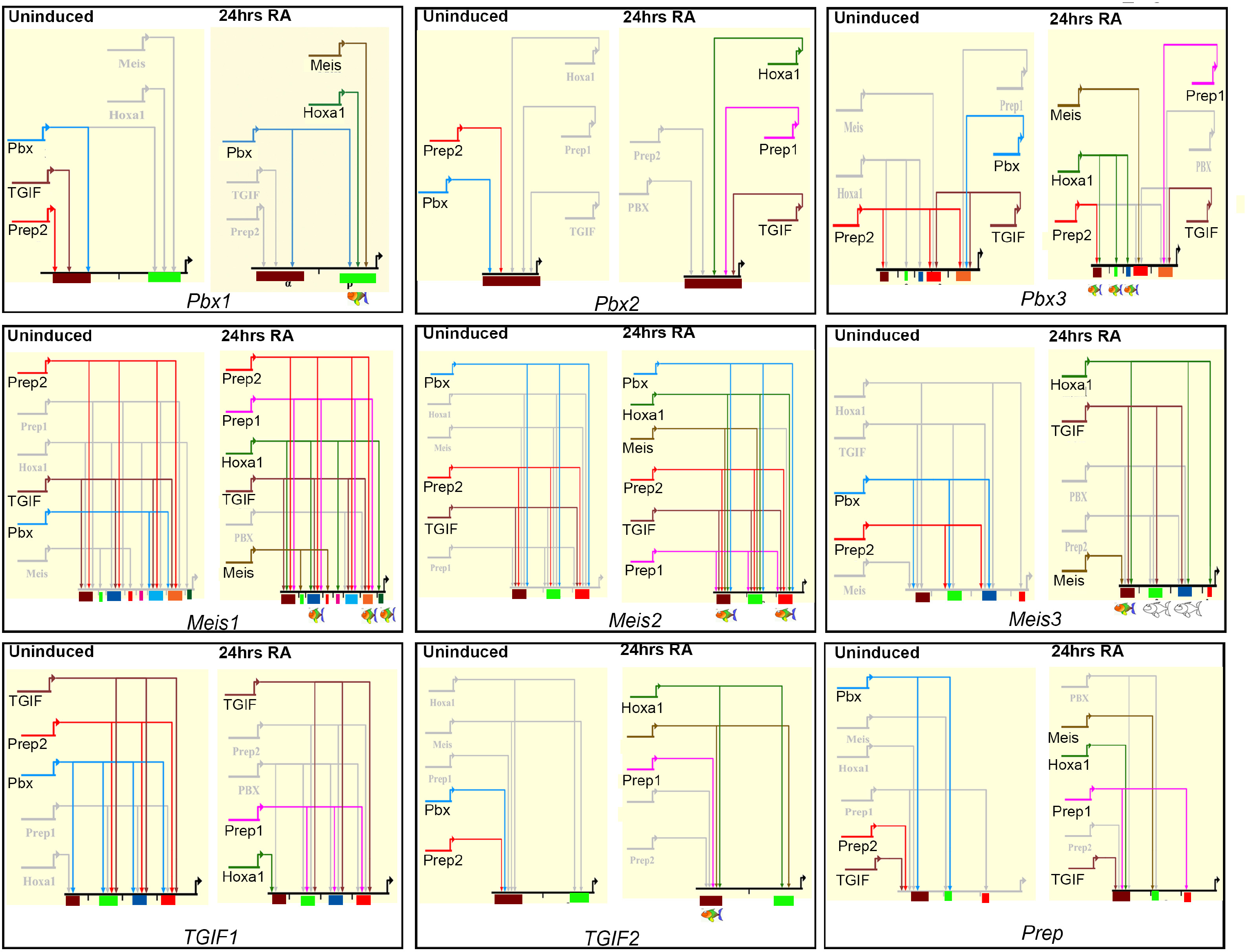
Biotapestry models depicting the auto- and cross-regulatory interactions between Hoxa1 and TALE members. In each plot ES cells are depicted on the left and differentiated cells on the right. The solid colored lines indicate binding of each specific TALE factor or Hoxa1, while grey lines indicate lack of binding in the given cellular states. At the bottom the colored boxes indicate each TALE locus and the respective cis-regulatory regions that integrate the specific binding inputs. In some cases, immediately adjacent binding regions are collapsed into a single cis-regulatory region and shown as one color. All cis-regulatory elements with a fish at the bottom were tested for regulatory potential in zebrafish transgenic reporter assays. Colored cartoon fish indicates those with spatially-restricted regulatory activity while the white fish indicate a lack of reporter activity.

## Discussion

We have identified and characterized regions bound by Hoxa1 on a genome-wide basis in differentiating mouse ES cells. These analyses indicate that Hoxa1 binding is frequently associated with a wide variety of motif combinations, including consensus sites for Pbx and Meis. High resolution mapping (ChIP-nexus) of individual binding peaks revealed evidence for multiple sites of Hoxa1 binding on a series of distinct consensus motifs for Hox, Hox-Pbx, Pbx and Meis (Fig. 1E and 5). Pbx and Meis show physical occupancy on these regions (Fig. 1D), indicating that Hoxa1 may be recruited to some of these sites through interactions with Pbx and Meis as cofactors. We found that the strong association of Pbx and Meis with Hoxa1 occupancy, extends to other members of the TALE family (Prep1, Prep2, TGIF) as supported by their genome-wide occupancy and binding site preferences (Fig. 2). Hence, TALE proteins represent a wide repertoire of Hox cofactors.

An important observation of this study is that nearly all Hoxa1 bound regions are associated with occupancy of diverse combinations of TALE proteins. These binding patterns fall into eight distinct clusters based on differential occupancy of TALE proteins. Motif enrichment analysis reveals that at the level of *cis*-elements these clusters are composed of a unique collection of enriched classes of TALE and novel motif signatures. Hence, the binding patterns found in Clusters1-8 relate to the underlying patterns of motif enrichment as a consequence of the presence of these particular sequences. This provides a mechanistic basis via a combinatorial *cis*-binding code, for recruitment of different permutations of TALE proteins to the Hoxa1-bound regions. The combination of multiple binding preferences and clustering of motifs for both Hoxa1 and TALE proteins may add to robustness and contribute to specificity to these regions as Hox response elements. This underscores the important general role that the TALE family proteins have in broadly modulating the specificity and function of Hox proteins. Furthermore, these distinct binding clusters are likely to be biologically meaningful because KEGG analysis implies that the target genes associated with the different clusters are involved in different processes (Supplemental Fig. S4). Together this suggests that the patterns of TALE binding and motif and pathway enrichment are causally related to each other.

Our analysis reveals that a common feature of Hoxa1 bound regions in differentiated cells is the presence of multiple sites for Hoxa1 and TALE cofactors. Comparative analysis of differential binding properties of Hoxa1 in ES cells and differentiated cells revealed that the presence of clustered motifs results in higher occupancy (Fig. 1B and Supplemental Fig. S2). This is consistent with the idea that clusters of suboptimal or low affinity binding sites can play an important role in potentiating specificity and input of TFs and their cofactors through cooperativity (Crocker et al. 2015; Farley et al. 2015).

The binding properties of these classes have revealed novel interactions between different TALE family members and Hoxa1. Published studies have shown that proximity of clustered Hox-Pbx and Pbx-Meis/Prep sites each bind their respective heterodimers, but also facilitate the formation of ternary Hox-Pbx-Meis/Prep complexes to fine tune binding specificity and potentiate regulatory activity (Berthelsen et al. 1998; Ryoo et al. 1999; Ferretti et al. 2000; Ferretti et al. 2005). The combinatorial binding patterns defined by clustering of Hox and diverse TALE motifs in this study provides further opportunities for protein-protein interactions. This opens the possibility that other TALE proteins may also facilitate the formation of ternary complexes with Hox, in a manner analogous to Pbx and Meis.

From a regulatory perspective, our analyses have uncovered an extensive series of auto- and cross-regulatory interactions among *Hoxa1* and *TALE* genes (Fig. 4-7 and Supplemental Fig. S5). We found that Hoxa1 can bind to *cis*-regulatory regions of all of the TALE genes and demonstrated that several function as Hox-responsive regulatory regions (e.g. *Meis2* & *Meis3*). Similarly, each TALE protein shows occupancy of regions around *Hoxa1* and other TALE genes, suggesting extensive cross-regulatory input into each other’s expression. We also uncovered evidence for wide-spread auto-regulatory mechanisms through occupancy of Hoxa1 and TALE proteins on their own *cis*- regulatory regions. This suggests a model (summarized in Fig 6B) whereby extensive auto- and cross-regulatory feedback interactions among *Hoxa1* and *TALE* genes results in regulation of *Hoxa1* and cofactors important for potentiating the downstream roles of Hoxa1 in patterning and development.

## Methods

### Generation of epitope tagged Hoxa1 cell line

Construct with Hoxa1-triple Flag-Myc were generated in pBS31 using recombineering approach. KH2 ES cells were engineered through lipofection and Hoxa1-triple flag-Myc were inserted in Col II locus and put under control of doxycycline inducible promoter. All cell lines were tested for karyotype stability. FACS Calibur was used for analysis of DNA content and to get indirect inference of karyotype stability. Cells were induced with RA and doxycycline and induction was confirmed through western hybridization.

### ChIP and ChIP-Nexus

Epitope tagged Hoxa1 KH2 ES cells were differentiated with doxycycline and retinoic acid for 24 hours. ChIP was done according to the Upstate protocol described in (Smith et al. 2010) with certain modifications. Cells were fixed by adding formaldehyde to media at a final concentration of 1% and by incubating at 37^o^C for 11 min and stopped by 1/10 volume of 1.25 M glycine with incubation at room temperature for 5 min. Cells were sonicated for 25 min in Bioruptor at high setting and 30 sec on-off cycle. Anti-Flag M2 antibody (Sigma, F1804) with sepharose A beads were used for IP. For ChIP nexus, Anti-Flag M2 antibody (Sigma, F1804) was attached with Dynabeads (200 μl per reaction) and the protocol performed as described by He Q and coworkers (He et al. 2015).

ChIP-seq and ChIP-nexus libraries were sequenced on the Illumina HiSeq 2500, 51bp single-end. Raw reads were aligned to the UCSC mm10 mouse genome with bowtie2 2.2.0 (Langmead and Salzberg 2012). Primary reads from each bam were normalized to reads-per-million and bigWig tracks visualized at the UCSC genome browser (https://genome.ucsc.edu/). Peaks were called with MACS2 2.1.0.20140616 (Zhang et al. 2008), parameters “-g mm -p 0.25 ‒m 5 50”. From each replicate, top 100,000 peaks based on p-value were compared with IDR 1.7.0 for reproducibility (https://sites.google.com/site/anshulkundaie/proiects/idr) and valid pairs with IDR p-value ≤ 0.01 were taken as the peak list. For ChIP-nexus, alignments were separated by strand, and peaks were called on strandwise BAM files using MACS2, parameters “-g mm –nomodel”, and no input. Peak summits were used for motif analysis using HOMER.

### Analysis of motif content

For Fig. 5 (A), motif content Hoxa1 bound regions were analyzed using the vertebrate motif set from Transfac v2016.2 (Matys et al. 2006) and searched with the FIMO program (Grant et al. 2011). Each peak set was given a background of 10 random coordinate sets, each with the same length and chromosomal distribution as the original peaks. Motif enrichment in peaks versus background was assessed with Fisher’s Exact Test, using BH p-value correction. To identify enriched known motifs in Hoxa1 and TALE proteins bound region and Hoxa1 bound region identified using ChIP-nexus, HOMER software package were used with parameter ‒size 100 (Heinz et al. 2010). Motif signature for TALE protein were generated using MEME suite. Enriched motifs were identified from genomic regions bound by individual TALE motifs using MEME (Bailey and Elkan 1994). A list of motifs was generated through combining all novel and consensus TALE motifs identified from genome-wide binding analysis of individual TALE factors. Redundant motifs with more than 60% similarity were identified using MAST (Bailey and Gribskov 1998). A final non-redundant motif list was generated by including the longest of these similar motifs and discarding the shorter variants. Nonredundant motifs were tested for their enrichment using AME (Buske et al. 2010) in individual TALE protein bound regions and a heatmap was generated through hierarchical clustering using p-values.

### Coverage heatmaps

Signal across peak coordinates was visualized with the CoverageView package in R (https://www.bioconductor.org/packages/release/bioc/html/CoverageView.html), using windows ± 5kb from peak midpoints. To focus sensitively on binding trend and not magnitude, IP z-score values were used for transcription factor samples. For ATAC-seq, we transformed the raw IP counts matrix to percent-of-maximum-value, so that values would be within the same general range as z-scores.

### Clustering

Coverage heatmaps for Hoxa1 and all TALE proteins, at Hoxa1 peaks, was generated by kmeans-clustered in R, using k from 2 to 20. A k=8 was chosen as this produced the best cluster validity metrics.

### Generation of reporter constructs

PCR-purified genomic regions were cloned into the HLC vector(Parker et al. 2014) using the Gibson Assembly Master Mix (NEB). Specific mutations to enhancer sequences were generated using Gibson assembly approach.

### Zebrafish transgenic lines

The following pre-existing zebrafish lines were used for experiments: Slusarski AB - wild type; *tg (mmHoxb1: EGFP*) - GFP expressed in r4 by the mouse *Hoxb1* auto-regulatory element (Parker et al, 2014); *egr2b:KalTA4BI-1xUASkCherry* ‒ mCherry inserted in the endogenous *egr2b* locus and expressed in r3 and r5(Distel et al. 2009).

### Zebrafish reporter assay

Zebrafish transgenesis was performed as previously described (Fisher et al. 2006). At least 100 embryos were injected for each construct. Embryos were screened for GFP reporter expression at approximately 24 hpf and 48 hpf with a Leica M205FA microscope and images were captured for fluorescent and brightfield signals with a Leica DFC360FX camera using LAS AF imaging software. Images were cropped and altered for brightness and contrast using Adobe Photoshop CS6. For certain constructs, transgenic lines were generated by selecting 25 GFP-expressing F0 embryos for raising to adulthood and outcrossing with wild-type fish, with the resulting F_1_ progeny being screened for GFP fluorescence indicative of germline transgene integration. GFP expression domains were considered consistent for a given enhancer if shared between three independent F1 transgenics.

## Acknowledgements

We thank Tari Parmely and members of the Stowers Institute (SIMR) Tissue Culture Facility for assistance with mouse ES cells and other cell culture needs, the SIMR Molecular Biology Facility for genomics and sequencing approaches, the SIMR Aquatics Facility for zebrafish care, Heidi Monnin and the SIMR Laboratory Animal Services Facility for mouse care, and members of the Krumlauf and Zeitlinger labs for valuable discussions and feedback. This work was performed to fulfill, in part, requirements for IRP Ph.D. thesis research as a student registered in the Stowers Institute Graduate Program.

## Additional information

### Competing interests

The authors declare that no competing interests exist.

### Ethics and Animal experiments

All experiments involving zebrafish (Protocol ID: 2015-0149) and mice (Protocol ID: 2016-0164) were done under approved protocols issued to REK as the PI by the Institutional Animal Care and Use Committee of the Stowers Institute for Medical Research.

### Funding

This research was supported by funds from the Stowers Institute to REK (Grant 1001).

### DATA submission

All raw sequencing data is submitted in NCBI as SRA accession number-SRP079975 and SRAPRJNA341679. All other original source data have been deposited in the Stowers Institute Original Data Repository and are available online at http://odr.stowers.org/websimr/.

### Author contributions

BDK, HJP, MEP, BDS, JRU and REK, experimental conception and design; BDK, HJP, MEP, IRP,BDS, and JRU, Acquisition of data; BDK, HUP, AP, IRP,BDS, JRU, JBZ, REK, Analysis and interpretation of data; BDK, HUP, MEP, AP,IRP, BDS, JRU, JBZ and REK, drafting and revising the article; MEP and JBZ, contributed unpublished data, protocols and reagents.

### Author information

Corresponding author and requests for materials should be sent to REK

Robb Krumlauf ORCHID ID http://ordd.org/0000-0001-9102-7927

